# Effect of Cellular and ECM Aging on Human iPSC-derived Cardiomyocyte Performance, Maturity and Senescence

**DOI:** 10.1101/2020.09.28.316950

**Authors:** S. Gulberk Ozcebe, Gokhan Bahcecioglu, Xiaoshan S. Yue, Pinar Zorlutuna

## Abstract

Cardiovascular diseases are the leading cause of death worldwide and their occurrence is highly associated with age. However, lack of knowledge in cardiac tissue aging is a major roadblock in devising novel therapies. Here, we studied the effects of cell and cardiac extracellular matrix (ECM) aging on the induced pluripotent stem cell (iPSC)-derived cardiomyocyte cell state, function, as well as response to myocardial infarction (MI)-mimicking stress conditions *in vitro*. Within 3-weeks, young ECM promoted proliferation and drug responsiveness in young cells, and induced cell cycle re-entry, and protection against stress in the aged cells. Adult ECM improved cardiac function, while aged ECM accelerated the aging phenotype, and impaired cardiac function and stress defense machinery of the cells. In summary, we have gained a comprehensive understanding of cardiac aging and highlighted the importance of cell-ECM interactions. This study is the first to investigate the individual effects of cellular and environmental aging and identify the biochemical changes that occur upon cardiac aging.

## 1. Introduction

Cardiovascular diseases (CVDs) account for one-third of the deaths worldwide[1]. Of the CVDs, myocardial infarction (MI) is the leading cause of global health loss and its occurrence is highly associated with age[2,3]. The prevalence of MI is only 0.6% among patients under 40, while it is 26.2% among patients over 80[1,2]. In addition, post-MI healing and repair of the damaged cardiac tissue in older patients are harder and it significantly lowers their life expectancy. The long term mortality rate among older patients over 75 years is 77%, while it is 45% for 65 to 74 years old patients [4]. Therefore not only the occurrence but also the post-MI cardiac recovery is highly influenced by age[4,5]. The age-associated increase in post-MI result severity might be due to the aged cells’ inability to respond to external factors such as cytokines[6], diminished regenerative capability of the cells, as well as the deregulations in the cardiac extracellular matrix (ECM) at advanced ages[7].

There has been extensive effort to develop cell-based therapies to treat MI patients. Stem cell therapy, the predominant approach used to treat MI, involves injection of stem cells to the infarct site with the aim of replenishing the aging cell population and regenerating the heart tissue. However, there is no clear evidence showing noticeable functional improvement in patients with CVD after stem cell injections[8], probably because of the loss of replicative abilities and the limited survival of these cells in the body, as well as the risk of differentiation to non-myocytes within the injured tissue[9]. Additionally, there is evidence for an inverse relationship between patient age and treatment success[6,10]. Alternatively, injection of lab-grown stem cell-derived cardiomyocytes (CMs) have shown partial integration with the host tissue, attenuation in adverse cardiac remodeling, and mechanical and functional cardiac improvements[11–13]. However, these cells remain mostly at the fetal stage and hold possible arrhythmia and tumorigenicity risks when used as therapeutics[14–17]. The main reason for the failure to obtain mature CMs *in vitro* is the inability to mimic the natural cardiac environment that provides biochemical and mechanical cues necessary for the complete differentiation of stem cells to CMs [18].

As a major component of the cardiac environment, extracellular matrix (ECM) is composed of both soluble and structural proteins necessary to maintain cardiac homeostasis [19,20]. Soluble factors, which include cytokines and other growth factors, have shown improvement in cardiac function when used in conjunction with stem cells [6,9,21–25]. Structural proteins such as collagen and hyaluronic acid have also been shown to improve cardiac function[26,27]. Hence, decellularized ECM, which contains both the soluble and structural proteins, has gained interest in the cardiac engineering field for its potential to repair the infarcted heart [28–33]. Injecting decellularized cardiac ECM of the neonatal mouse was shown to support post-MI regeneration and reduce MI-induced fibrosis in adult mouse models [34]. Although adult and neonatal cardiac ECM biochemical compositions are often compared to explain such study outcomes, little is known about the cardiac ECM at older ages and how it interacts with CMs. ECM undergoes constant remodeling to adapt to changing conditions, including disease and aging [35], which clearly makes the ECM at young ages distinct from that at older ages or after MI. Thus, the interplay between cells and ECMs at various ages remains to be studied which will be essential for developing novel cell or ECM-based therapies [7,36].

The purpose of this study was to investigate the individual effects of cellular and ECM aging on cardiac maturity and function, as well as the cells’ response to MI-like stress conditions. To this end, we chronologically aged human induced pluripotent stem cell (iPSC)-derived cardiomyocytes (iCMs) based on our previous study where we had demonstrated that cells went through accelerated senescence and showed functional deterioration within 4 months in culture, resembling 65-year-old human cardiac tissue [37]. We seeded these aged cells and their controls (1-2-month-old young cells) on decellularized heart ECMs from mice of three age groups: (i) 1-3 month, (ii) 6-9 months and (iii) 22-24 months, to recapitulate ECMs of young (10-16-year-old), adult (24-32-year-old) and aged (67-73-year-old) humans[38–40], respectively. We investigated the proliferation ability, senescence state, and the cardiac maturity and function-related electrophysiology of the young and aged iCMs on the young, adult, and aged cardiac ECM-coated glass surfaces. Furthermore, we performed proteome profiling of the iCMs and studied the changes in response to the different age ECM environments, as well as to the MI-mimicking stress conditions. We finally assessed the cellular response to MI-mimicking conditions.

To the best of our knowledge, this is the first study to investigate the individual effects of cellular and environmental aging on cell behavior and to identify the biochemical changes that occur in the cardiac tissue microenvironment upon aging. Given the complexity of the heart, dissecting the individual effects of cellular and environmental aging would be of utmost importance for advancing our understanding of cardiac aging, and for developing effective treatment strategies for heart diseases based on patient’s age.

## 2. Materials and Methods

### 2.1. Mouse heart isolation and handling

All animal procedures were performed in accordance with the Institutional Animal Care and Use Committee (IACUC) at the University of Notre Dame. Hearts from C57BL/6J mice (000664/Black 6, female) of three age groups, 1-3 months, 6-9 months, and 22-24 months, were isolated to study young, adult, and aged cardiac microenvironments, respectively. Tissues were either flash frozen and stored at -80°C until further use or immediately embedded in the Optimal Cutting Temperature compound (OCT, VWR) and stored at -20°C until immunohistochemical staining. Flash-frozen tissues were equilibrated to room temperature (RT), washed in phosphate buffered saline (PBS, VWR) and were cut into smaller pieces prior decellularization. The OCT embedded tissues both in native and decellularized forms were processed at the Notre Dame Integrated Imaging Facility and stained for hematoxylin and eosin and Masson’s trichrome.

### 2.2. Heart decellularization and characterization

The mouse hearts were first decellularized in 0.25% and 0.5% (wt/vol) sodium dodecyl sulfate (SDS, VWR) solution at RT for 24 h or until the tissues turn transparent white with constant agitation. Then, samples were transferred to the Triton X-100 (Sigma-Aldrich) solution at the same concentration as the respective SDS solutions for 30 min with constant agitation. Decellularization was performed with low concentrations of both detergents to achieve maximum decellularization with minimum disruption of the tissue composition in a non-perfusing setup. Once the decellularization is done, samples were washed thoroughly with DI water to remove any residual detergent.

Mass spectrometry (MS) was used to determine the protein composition of the decellularized mouse heart tissues (n=3-4 for each developmental age). The samples were digested in the digest solution (5M urea, 2M thiourea, 100mM Ammonium bicarbonate and 50mM Dithiothreitol, pH 8.0) at 4°C for 24 h with constant stirring. The resulting soluble and insoluble ECM components were processed separately as the supernatant via in-solution trypsin digestion [97] and the pellet via sequential CNBr and trypsin in-gel digestion protocols [98]. The protein concentration of the samples was measured with the Pierce Rapid Gold BCA Assay (Thermo Scientific) and equalized prior to MS. The most abundant ECM components for each developmental age were identified.

### 2.3. Cardiac ECM solubilization and characterization

Decellularized ECMs were lyophilized and pulverized with liquid nitrogen. ECM powder was digested in a 1 mg/mL pepsin (Sigma-Aldrich) in 0.1M HCl (10:1, w/w, dry ECM:pepsin) at RT with constant stirring until a homogeneous solution was obtained. The insoluble remnants were removed by centrifugation, the supernatant was neutralized using 1M NaOH solution, and used immediately to prevent degradation. Prior to experiments, we measured the total protein concentrations using Rapid Gold BCA Assay (Thermo Scientific) and diluted ECM solutions to 1mg/ml with Phosphate-Buffered Saline (PBS, Corning).

### 2.4. iPSC cardiac differentiation and cell culture

DiPS 1016 SevA from human dermal fibroblasts (Harvard Stem Cell Institute iPS Core Facility) were used for cardiomyocyte differentiation. The hiPSCs were maintained and differentiated using previously established protocols [37,41,99]. Briefly, hiPSCs were cultured on Geltrex (1%, Invitrogen)-coated tissue culture flasks in mTeSR-1 media (StemCell Technologies). At 80-85% confluency, the hiPSCs were collected and seeded at 1.5 × 10^5^ cells/cm^2^ cell density on glass coverslips with Rho-associated coiled-coil containing protein kinase (ROCK) inhibitor supplemented mTeSR media (5 *μ*M, StemCell Technologies). The culture was maintained with daily media changes until 90% confluency was reached. The cardiac differentiation was initiated using the canonical Wnt pathway. First, hiPSCs were treated with the RPMI 1640 (Life Technologies) media supplemented with B27 without insulin (2%, Invitrogen), β-mercaptoethanol (3.4 × 10^−4^%, Promega) and Penicillin (1%) (CM(−)) with the addition of Wnt activator CHIR99021 (12 *μ*M, CHIR, Stemgent) (Fig. 1A). On day 2, media was changed to CM(−), on day 4, cells were treated with Wnt inhibitor IWP4 supplemented CM(−) (5 µM, Stemgent) and on day 6, media was changed back to CM(−). From day 9 on, cells were maintained in RPMI 1640 media supplemented with B27 with insulin (2%, Invitrogen), β-mercaptoethanol and Penicillin (CM(+)) media with a media change every 3 days. iCMs were cultured for 35-55 days and used as young cells or for 100-120 days and used as aged cells [37].

**Figure 1.**
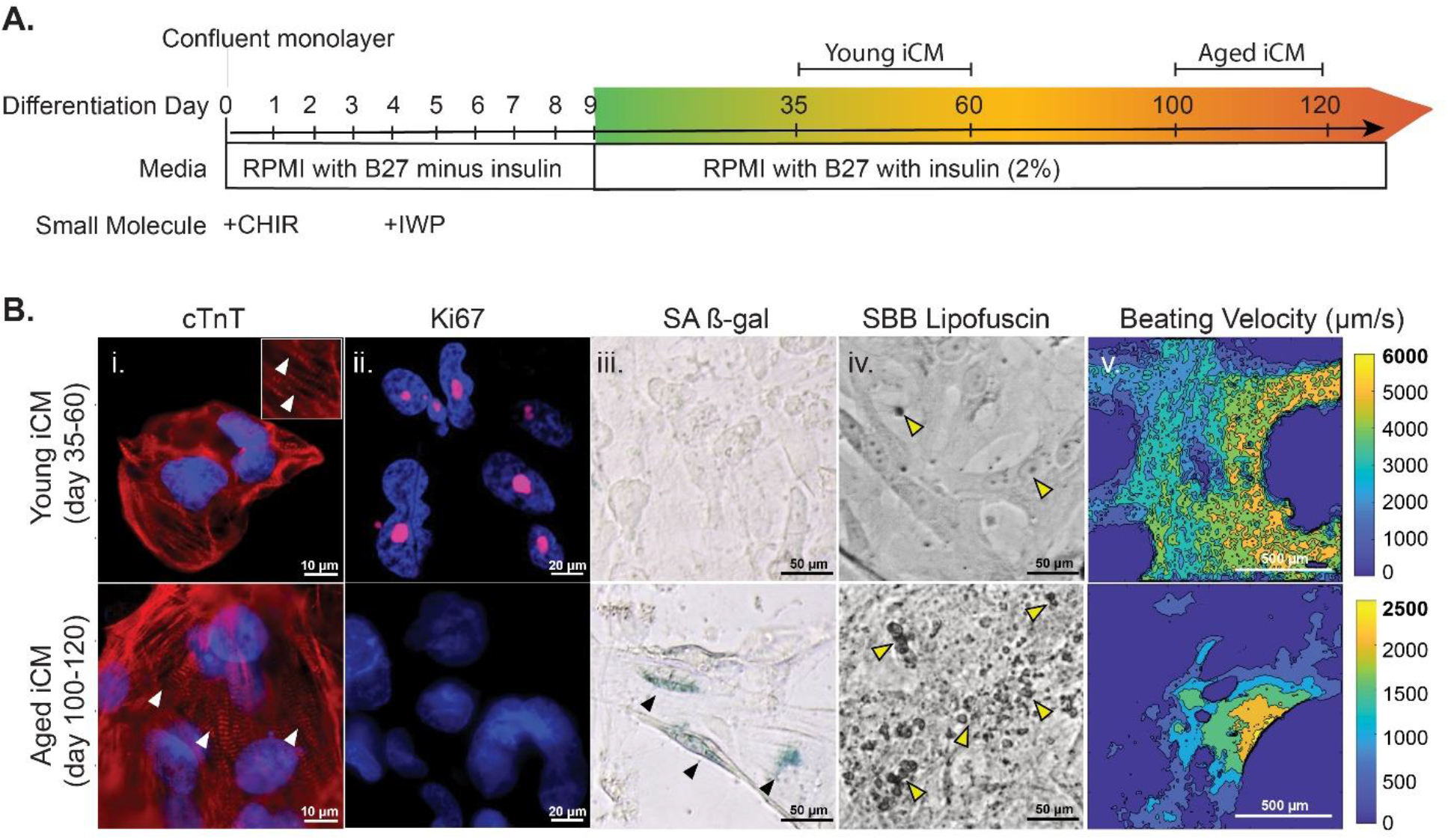
Characterization of iCMs after regular and prolonged culture. (A) Differentiation protocol for generating cardiomyocytes from iPSCs. Cells were cultured for either 35-60 days (young iCM) or 100-120 days (aged iCM). (B) Comparison of young and aged cells for (i) cTnT (red), (ii) proliferation marker Ki67 (magenta), senescence markers (iii) SA β-gal (blue) and (iv) lipofuscin (black), and (v) spontaneous beating velocities of the cell sheets. Nuclei were counterstained with DAPI (blue).

### 2.5. ECM coating and iCM seeding

Glass coverslips were coated with solubilized ECM at 37 °C for 30 min. Fibronectin (50μg/ml, Sigma-Aldrich) was used as the control coating protein. The iCMs were trypsinized and seeded at 2.5 × 10^5^ cells/cm^2^ cell density on ECM-coated glass coverslips with Dulbecco’s Modified Eagle Medium (DMEM, Hyclone) supplemented with 5% FBS. Cells were maintained in CM(+) media for 21 days with a media change every 3 days.

### 2.6. Senescence associated characterization of iCMs

Age-related phenotype was determined using senescence-associated β-galactosidase assay (SA-β-gal, Millipore) and Sudan Black B (SBB, Sigma-Aldrich) staining for lipofuscin as previously described [100,101]. After a thorough wash with PBS, cells were imaged under a bright-field microscope (Leica). Lipofuscin accumulation was quantified as the area covered by lipofuscin droplets using ImageJ software.

### 2.7. Electrophysiological characterization of iCMs, and drug response

As an indicator of cardiac function, contractile properties of the cells were studied as previously described[37]. Briefly, three representative regions from each sample were selected and spontaneous beatings were recorded at a rate of 30 frames per second for 20 seconds. Using a custom-developed MATLAB code, each video frame was divided into analysis regions (16×16 px) and with the block matching algorithm, velocity vectors were generated, and maximum displacement, and motion velocity and direction were calculated. Heat maps of the beating iCMs were generated to determine the general beating trend of the cells.

The calcium (Ca^2+^) transient of iCMs was assessed to further investigate the electrophysiology. Cells in culture washed with PBS and the media was replaced by Ca^2+^-sensitive Fluo-4 AM (Life Technologies) solution, as instructed by the manufacturer. Cell beating was recorded real-time using a fluorescence microscope (Axio Observer.Z1, Zeiss, Hamatsu <u>C11440</u> digital camera) at 30 ms exposure for 30 sec. To assess the drug response, we treated cells with isoproterenol (1 µM, ISO) for 10 min at 37°C. Spontaneous beating before and after ISO was recorded. The baseline for calcium transient and an initial beat rate of the cells were obtained from pre-drug recordings (5 images/well, and 3 wells/age group). The beating rate after ISO treatment relative to that before treatment was calculated by dividing post-ISO beating by pre-ISO beating. Rate of Ca^2+^ release (time to peak Ca^2+^ transient amplitude) and action potential duration at 50% (APD50) and 90% (APD90) of the amplitude was calculated from the raw data obtained from intensity versus time plots.

### 2.8. I/RI experiment

RPMI 1640 no glucose media (Life Technologies) supplemented with B27 minus antioxidants (2%, Invitrogen), β-mercaptoethanol (3.4 × 10^−4^%, Promega) and Penicillin (1%) (AO(−)) was equilibrated to the hypoxia (1% oxygen) for overnight. To mimic ischemia (I), iCMs were treated with the hypoxia-equilibrated AO(−) media and incubated in the hypoxia chamber set to 1% oxygen for 3 h. To mimic reperfusion injury (RI), iCM media was changed to RPMI 1640 media supplemented with B27 minus antioxidants (2%, Invitrogen), β-mercaptoethanol (3.4 × 10^−4^%, Promega) and Penicillin (1%) and iCMs were incubated in the normoxia (21% oxygen) for 12 h.

### 2.9. Post I/RI analysis

At 12 h RI, we stained cells with Live/Dead assay (Life Technologies) and calculated survival as the ratio of live cell number (subtracting dead cell number from total) to total cell number. At 3 h RI, ROS levels in the culture were determined via Mitochondrial ROS Activity Assay Kit (eEnzyme). Briefly, we prepared the stain solution as instructed, treated cells for 30 min at 37°C and monitored fluorescence signal immediately after under a fluorescent microscope. Post I/RI ROS generation was normalized to the total cell number.

### 2.10. Proteome Arrays

The relative cytokine content of the decellularized mouse heart tissues (n=3) were obtained using the Mouse XL Cytokine Array Kit (ARY028, R&D Systems). Briefly, array membranes were blocked with an array buffer for 1h at RT. Tissue extracts were obtained by 1% Triton-X incubation, diluted and incubated overnight with the membranes at 4°C. The unbound proteins were rinsed away, the membranes were incubated with the biotinylated antibody cocktail.

The relative expression levels of cellular stress related proteins of iCMs (n=3) were analyzed using the Proteome Profiler Human Cell Stress Array (ARY018, R&D Systems), according to the manufacturer’s instructions. Briefly, adherent cells were lysed, and incubated with the detection antibody cocktail for 1h at RT. Meanwhile, the array membranes were blocked with an array buffer for 1h at RT. The sample/antibody mixture was incubated overnight with the array membranes at 4°C. The relative expression levels of apoptosis related proteins of post I/RI iCMs (n=3) were analyzed using the Proteome Profiler Human Apoptosis Array (ARY009, R&D Systems), according to the manufacturer’s instructions. Briefly, adherent cells were lysed, and the array membranes were blocked with an array buffer for 1h at RT. The cell lysates were diluted and incubated overnight with the membranes at 4°C. The unbound proteins were rinsed away, the membranes were incubated with the biotinylated antibody cocktail.

For all, protein concentrations were normalized before starting the arrays. At the end, the unbound proteins were rinsed away, the membranes were incubated with the Streptavidin-HRP, then developed using chemiluminescent detection reagent mixture. For quantification, the background was removed, and the pixel density of each spot was measured using ImageJ.

### 2.11. Immunostaining

Immunostaining was performed on the adherent monolayer cell culture. The cells were washed with PBS, fixed with paraformaldehyde (4% PFA) for 15 min at RT and treated with the permeabilization solution (0.1% Triton-X 100) for 20 min at RT. Cells were then treated with the blocking solution (10% Goat serum) for 1 h at RT and incubated with the primary antibodies for TNNT2 (Abcam, ab45932), Ki67 (Thermo Fisher, 14-5698-82), cytochrome c (Novus Biologicals, 7H8.2C12), cleaved caspase-3 (R&D System, MAB835) (1:150 in goat serum) overnight at 4 °C. Cells were then washed with PBS and incubated with the species-appropriate secondary antibodies (Life Technologies) (1:300) for 6 h at 4 °C with a thorough PBS wash after each incubation. Images of cells were captured using a fluorescence microscope (Zeiss, Hamamatsu ORCA flash 4.0) and Post imaging processing was performed using Zeiss Zen software (fluorescence microscope). The cardiac phenotype was analyzed primarily via immunostaining for a cardiac marker TNNT2 and the sarcomere lengths were measured via the ImageJ software (NIH) (n=10 per ROI).

### 2.12. Statistical Analysis

The mean ± standard deviation (SD) was reported for all replicates. 2-way analysis of variance (ANOVA) was used to test the interaction between the cell age and the ECM age. In case of interaction, the data were re-analyzed for each cell age group. One-way ANOVA was applied to test the effect of ECM groups on young and aged cells separately. One-way ANOVA with post-hoc Tukey’s test was used to assess the statistically significant differences using IBM SPSS Statistics Software version 25. All *p* values reported were two-tailed, and *p* < 0.05 was considered statistically significant. Sample size (n) ≥ 4 for individual experiments. The Pearson correlation test was used to calculate the correlation between ECM age and Lipofuscin accumulation. Pearson correlation coefficient (r) is calculated as:

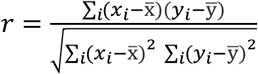

## 3. Results

### 3.1. Prolonged culture mimics chronological aging

Human iPSCs (DiPS 1016 SevA) were differentiated into CMs using the canonical Wnt pathway [41]. Following our previously established protocol [37], we aged our iCMs chronologically by prolonging the culture period. We cultured the iCMs for either 35-55 days to generate young cells or 100-120 days to mimic aged cells **(Fig. 1A)**. To verify the effect of prolonged culture on cardiac maturity, cells were first stained for the cardiac marker tropomyosin-binding subunit of the troponin complex (cTnT). The majority of the differentiated cells marked positive for cTnT. The young iCMs were relatively small and round, whereas the aged iCMs were larger and rod-shaped with more aligned myofibrils **(Fig. 1Bi)**, indicating a more mature phenotype. Next, the cell senescent state was investigated through cells’ proliferative abilities and senescence marker expression. A larger fraction of the young cells expressed the proliferation marker Ki67; the number of aged cells presenting Ki67 was negligibly low (**Fig 1Bii**).

Conversely, only the aged cells displayed SA β-gal activity and lipofuscin accumulation (**Fig 1Biii-iv**). Furthermore, we analyzed cells’ spontaneous beating. The young cells showed robust and synchronized beating with almost 3 times higher maximum beating velocity than the aged cells displaying nonsynchronous beating (**Fig 1Bv**).

### 3.2. Decellularized ECM preserves characteristics of young, adult and aged hearts

Heart tissues were collected from 1-3-month (young), 6-9-month (adult) and 22-24-month-old mice (aged) corresponding to human ages of 10-16 years, 24-32 years, and 67-73 years, respectively **(Supplementary Fig. 1)** [38,40,42], and decellularized. Hematoxylin & Eosin and Masson’s Trichrome staining revealed successful decellularization (no nuclei were observed), and preservation of ECM protein structure **(Fig**.**2A)**. Collagen abundance (blue color in Trichrome staining) and the unit fiber thickness gradually increased with ECM aging.

**Figure 2.**
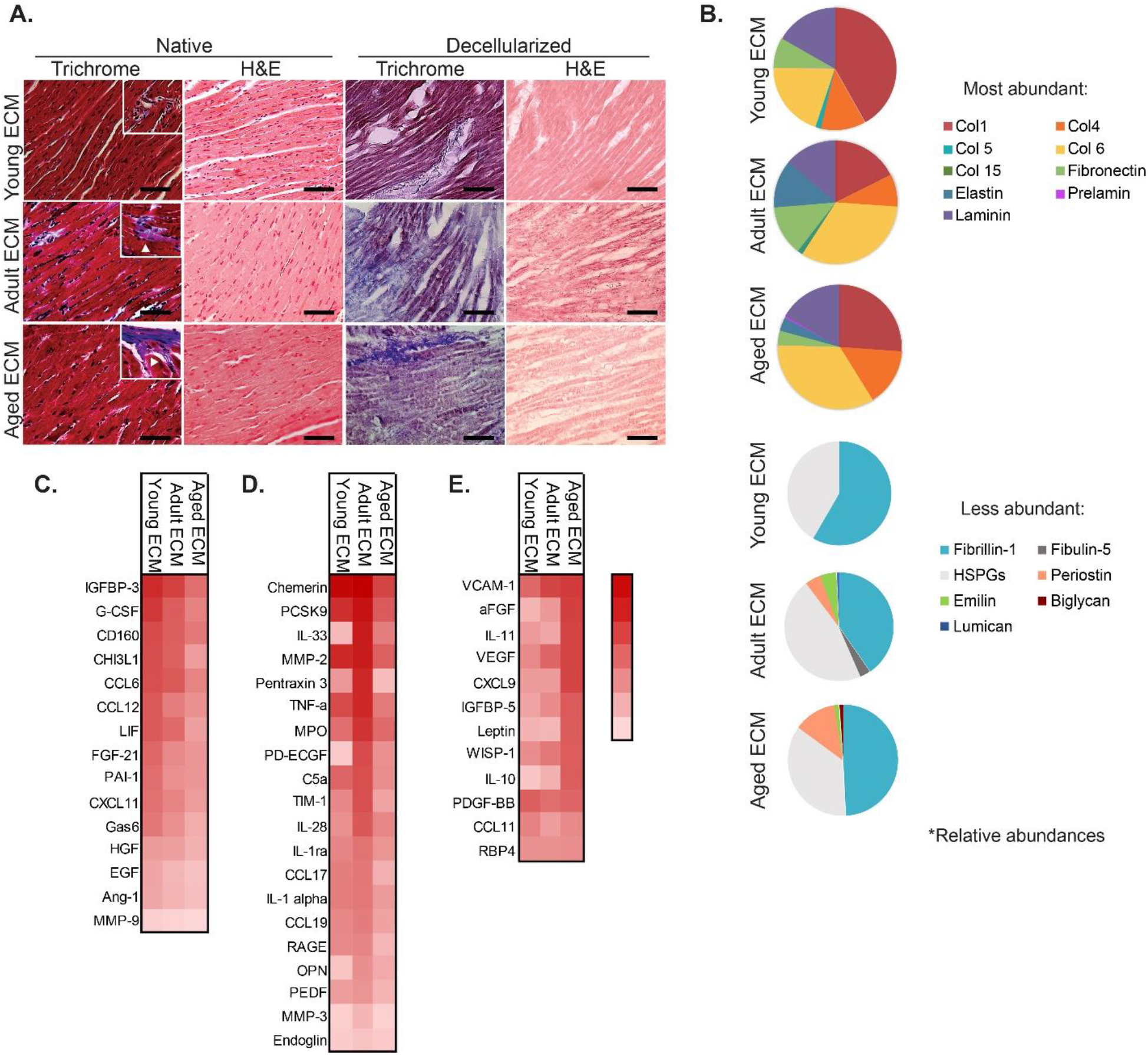
Biochemical characterization of ECM. (A) Histological analysis of native and decellularized cardiac tissue of young, adult, and aged mice. Scale bar represents 500 µm. (B) ECM compositions represented as the relative percentage of the most (top) and less (bottom) abundant proteins detected by mass spectroscopy. (C-E) Cytokines found in mouse heart ECM, split into subgroups showing the most abundant cytokines found in (C) young ECM, (D) adult ECM, and (E) aged ECM. Cytokines are listed in descending order within their respective groups.

Mass spectrometry analysis of the most abundant 15 ECM components revealed up to a 2-fold increase in some structural ECM proteins, including collagen I and IV, and laminin, in aged ECM compared to adult ECM **(Fig. 2B)**. Similarly, fibrillin-1 and periostin, which normally increase upon fibrosis- and aging-associated remodeling of the heart [43,44], were expressed to a higher extent in the aged ECM **(Fig. 2B)**. Contrarily, the elastic ECM proteins, fibronectin, elastin, as well as minor components, emilin-1 and fibulin-5 [45–49], were expressed to a lesser extent in the aged ECM (**Figs. 2B and 2C)**.

The biochemical composition of the decellularized ECMs was further assessed via cytokine antibody arrays measuring 111 different cytokines, chemokines, and other secreted entities. Among these factors, longevity, self-renewal, proliferation, and survival-associated factors, such as Fgf-21, G-Csf, Chi3l1, Lif, and Gas6 [50–55] were identified in the young ECM **(Fig. 2C)**. Cardioprotection, cardiac function, ECM remodeling, and antioxidation-associated factors, such as Pedf, Ptx3, Il-33, Pdecgf, and Opn [56–60]were present at high levels in the adult ECM **(Fig. 2D)**. Finally, fibrosis, post-MI cardiac remodeling, and hypertrophy associated factors such as Cxcl9, Il10, Il11, aFgf [61–64] were high in the aged ECM **(Fig. 2E)**. The above findings revealed notable alterations in the cardiac ECM biochemical composition upon aging and provided the rationale for our further studies investigating ECM age influence on cell state, behavior, and function. Decellularized ECMs were solubilized and used as coating materials to study cell-ECM interactions in young, adult, and aged cardiac microenvironment **(Fig. 3A)**.

**Figure 3.**
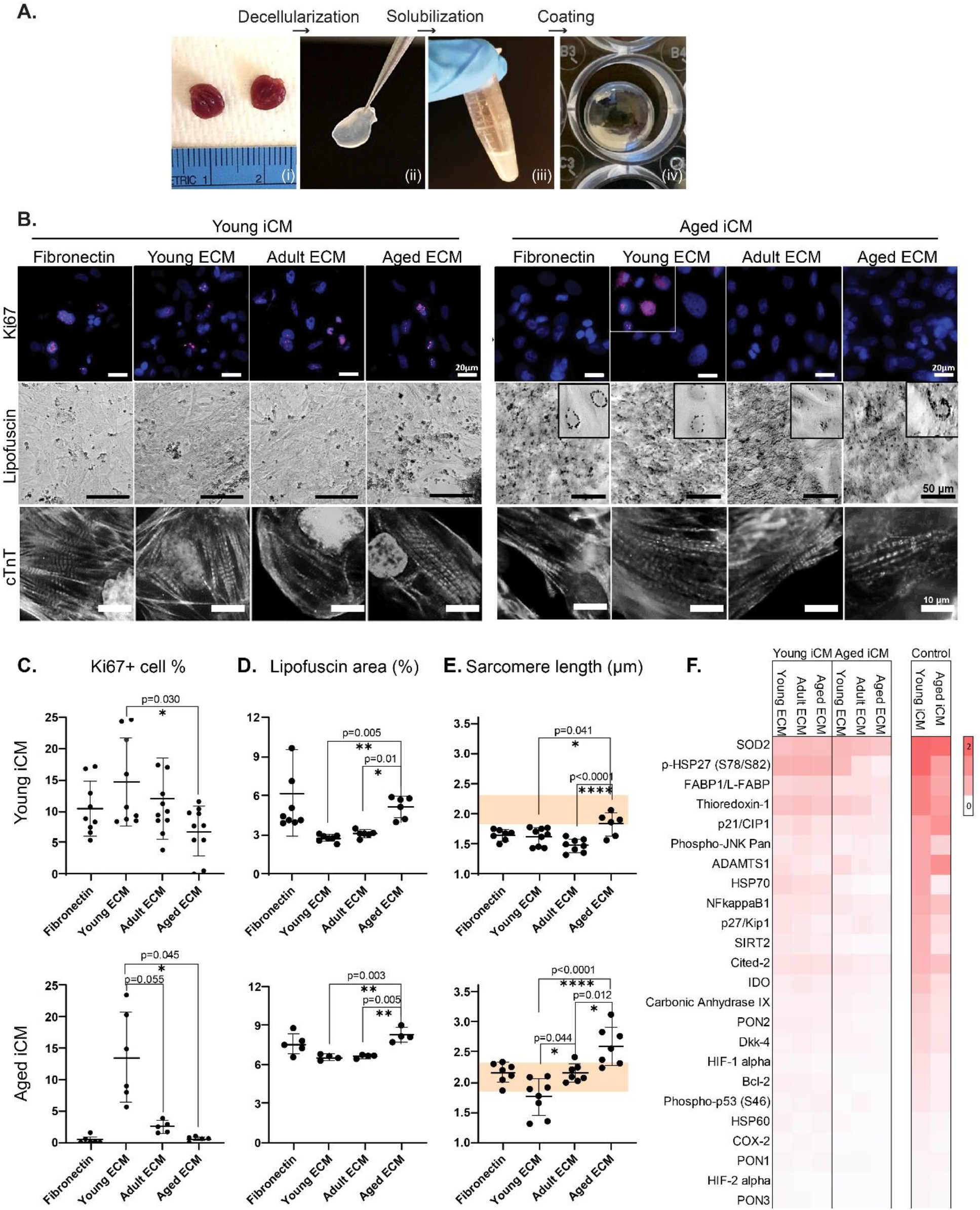
Cellular state assessment of iCMs. (A) Decellularization (i>ii), solubilization (ii>iii) and surface coating (iii>iv) protocol for mouse cardiac tissue. (B) Proliferation, senescence, and cardiac maturity related staining of the young (left) and aged (right) cells. (C) Proliferative cell percentage calculated as Ki67 presenting cell ratio. (D) Senescence-associated lipofuscin accumulation calculated as the covered area. (E) Sarcomere lengths measured from cTnT immunostaining of the cells. The shaded area indicates the adult human cardiomyocyte sarcomere length range [65]-[68]. (F) Performance baseline assessment of the cells through stress associated protein expressions. Statistical analysis was done using one-way ANOVA with post-hoc Tukey’s test. **** p<0.0001, ***p<0.001, **p<0.01, *p<0.05, n≥3. Data presented as mean ± standard deviation (SD).

### 3.3. Young ECM promotes proliferative abilities

Proliferation ability of the cells was assessed as the percentage of the Ki67 presenting cells. Our results showed an inverse relation between the proliferation rate and the ECM age. We observed less proliferative cells as the ECM age increased. Around 15% of the young cells were proliferative on the young ECM, while only around 7% on the aged ECM (p<0.05) **(Figs. 3B and 3C)**. Proliferative cell ratio significantly dropped with cell aging. On both adult and aged ECMs, a negligibly low amount of the aged cells was proliferative (<5%). More interestingly, the young ECM promoted aged cell proliferation and the proliferative aged cell ratio increased to ∼14% (p<0.05), which is comparable to of the young cells (∼15%) **(Fig. 3C)**. We also captured actively dividing aged cells on the young ECM **(Supplementary Fig. 2)**.

### 3.4. Aged ECM promotes aged phenotype

The senescent state with respect to ECM age was investigated via lipofuscin accumulation. We observed more lipofuscin accumulation with the increasing ECM age **(Figs. 3B and 3D)**. For both cell ages, the mean lipofuscin accumulation positively correlated with ECM aging (r=0.90 for young iCM, r=0.93 for aged iCM) **(Supplementary Fig. 3)** where lipofuscin covered area was significantly larger on the aged ECM than on the adult and young ECMs (p<0.01) **(Fig. 3D)**. Moreover, the accumulation rate was also positively correlated with chronological aging (r=0.85) The aged cells accumulated lipofuscin mainly in the perinuclear zone, and accumulation (the area occupied by lipofuscin) was halved in the young cells **(Figs. 3B and 3D)**.

### 3.5. Aged ECM elongates sarcomere length

The cardiac phenotype was assessed via the sarcomere length. The mean sarcomeric lengths of the young (1.47-1.83 μm) and aged (1.76-2.59 μm) cells were comparable to the fetal (1.6 μm) and adult (2.2 μm) human cardiomyocytes, respectively [65–68] (**Fig. 3E**). For the young cells, we measured similar sarcomere lengths on the young ECM (1.62 ± 0.14 μm) and adult ECM (1.47 ± 0.10 μm). On the aged ECM, the young cells had significantly elongated sarcomeres (1.83 ± 0.17 μm) (p<0.05). For the aged cells, sarcomere lengths elongated with increasing ECM age. We measured the shortest sarcomere length on the young ECM (1.76 ± 0.28 μm) and the longest on the aged ECM (2.60 ± 0.30 μm) (p<0.0001). Regardless of the cell age, we observed longer sarcomeres when cells were cultured on the aged ECM (p<0.01); young cells’ sarcomeres reached average lengths of a mature human CM, whereas the aged cells’ exceeded the upper limit of a healthy CM [66].

### 3.6. The cellular response on different age ECMs

The ECM effect on the cells was assessed in terms of the basal cell stress levels. In our control groups, we detected more pro-survival and antioxidant proteins such as SOD2, TXN, and HSPs [69–71] secreted from the young cells than the aged cells. Additionally, we detected higher ADAMTS1, an important mediator of cardiac aging, and p21 expression in the aged cells **(Fig. 3F)**. When ECMs were added to both young and aged cells, the stress related protein expressions were minimized. We did not observe any significant influence of the ECM age on cells’ basal stress levels. These results established a performance baseline for the cells cultured on different age environments that aid us to interpret the cell response we observed upon I/RI mimicking stress conditions.

### 3.7. Adult ECM supports cardiac function

The vector heat maps visualizing the spontaneous beating of the cells provided a comprehensive picture of the cardiac cell function (**Fig. 4A**). Young cells displayed homogeneous tissue-like beating on adult ECM and asymmetric beating with a center of contraction on young and aged ECM. Aged cells, on the other hand, showed a patchy beating pattern on all ECM age groups with larger beating areas on young and adult ECM.

**Figure 4.**
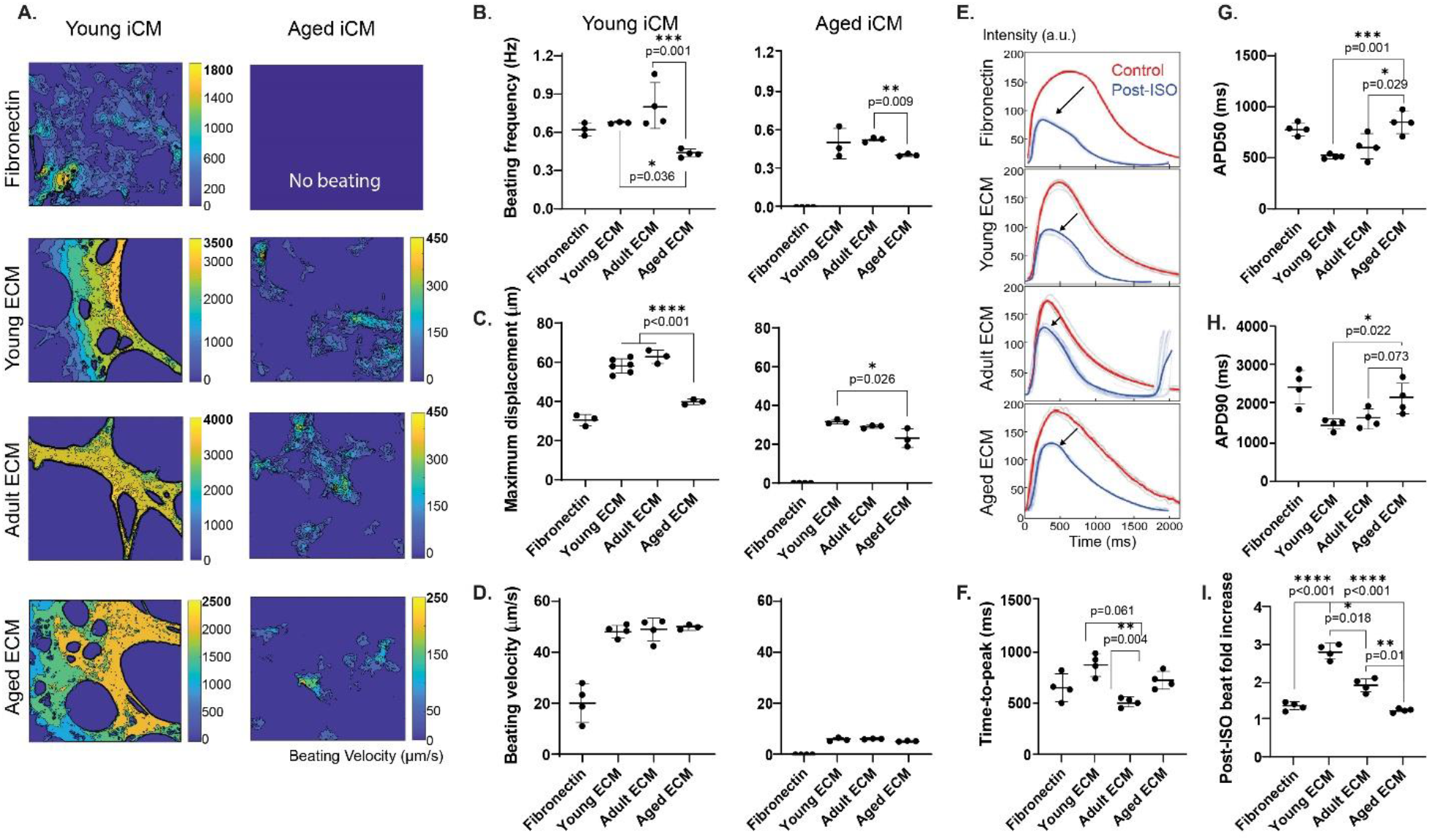
Mechanical and functional assessment of iCMs. (A) Heatmaps showing the beating velocity magnitude and distribution of young(left) and aged cells (right). Quantification of (B) beating frequency, (C) maximum displacement of a pixel in a frame due to spontaneous beating, and (D) beating velocity of spontaneous cell beating. (E) Action-potential curves of young cells before (control,red) and after (post-ISO, blue) isoproterenol treatment. Calculated (F) time-to-peak, (G) 50% decay, (H) 90% decay durations and (I) beat fold increase. Statistical analysis was done using one-way ANOVA with post-hoc Tukey’s test. **** p<0.0001, ***p<0.001, **p<0.01, *p<0.05, n≥3. Data presented as mean ± standard deviation (SD).

Detailed analysis revealed that the young cells performed significantly better, with faster, stronger, and more robust beating than the aged cells (**Fig. 4B-D**). Young cells beat at 0.43-0.80 Hz, whereas aged cells at 0.40-0.52 Hz, both exhibiting the lowest average beating frequency on the aged ECM **(Fig. 4B)**. Similarly, the maximum displacement was the lowest on the aged ECM, regardless of the age of cells **(Fig. 4C)**. The beating velocity, however, did not show any dependency on the ECM groups **(Fig. 4D)**. Overall, as in the aged heart, we observed a cardiac function deterioration with the aging of both cells and ECM. Young cells beat at a significantly higher frequency with a stronger pull on the young and adult ECMs compared to the aged ECM. Aged cells displayed a subtle response to ECM age, yet followed a similar trend to that of young cells.

### 3.8. Young and adult ECM improves calcium handling abilities

The calcium transient during spontaneous beating of cells was recorded and maximum contraction (time-to-peak, TTP), 50% decay (APD50), and 90% decay durations (APD90) were calculated. The aged cell beating was too weak, hence we only analyzed young cells for their calcium handling abilities. We observed a ventricular-type action-potential (AP) profile only for the young iCMs seeded on the adult ECM **(Fig. 4E)**. When quantified, cells displayed faster calcium transient on young and adult ECM where the smallest TTP was on the adult ECM, and the smallest basal APD50 and APD90 were on the young ECM **(Figs. 4F-H)**. Stimuli responsiveness was assessed via changes observed in beating profile upon isoproterenol (ISO) treatment. We observed a gradual decrease in drug responsiveness as the ECM age increased. Cell beat frequency increased almost 3-folds on the young ECM, 2-folds on the adult ECM, and very minimal on the aged ECM **(Fig. 4I)**. Overall, we observed enhanced calcium handling ability on the adult ECM and amplified drug responsiveness on the young ECM.

### 3.9. Aged cells respond poorly to MI-mimicking stress conditions

Myocardial infarction (MI) was simulated *in vitro* by culturing iCMs in hypoxia followed by normoxic media change. We incubated the cells in hypoxia-equilibrated glucose-deficient media as in the ischemic (I) tissue and made an immediate change to normoxic glucose bearing media to cause reperfusion injury (RI). Upon I/RI, we observed an overall decreased viability, increased reactive oxygen species (ROS) generation as well as activated apoptotic pathways compared to the normal cell culture conditions **(Fig. 5A)**. Post-I/RI cell viability was assessed via the live cell ratio in the culture. Survival of the young cells was the highest on the adult ECM (66.56 ± 7.85 %) compared to other ECM age groups, however, the aged cell survival did not show any dependency to the ECM groups (56.50± 12.22 % on young, 56.22± 8.74 % on adult, 56.40± 11.16 % on aged ECM) **(Fig. 5B)**. Because many of the dead cells detach or clump, especially in the aged iCM samples, the actual effect of these findings should be more pronounced than observed. In addition to survival analysis, activation of apoptotic pathways were investigated by ROS generation, cytochrome c (Cyt C) release into the cytosol, and activated caspase 3 (clvCas3) expressing cell ratio. We detected more ROS generated from aged cells, especially when cultured on aged ECM. Aged cell ROS generation increased with increasing ECM age; there were 4-folds more ROS on the aged ECM than on the young (p<0.0001) (**Fig. 5C**). Similar to ROS generation results, Cyt C release (p<0.01) (**Fig. 5D**) and clvCas3 activity (p<0.05) (**Fig. 5E**) were higher in the aged cells compared to young.

**Figure 5.**
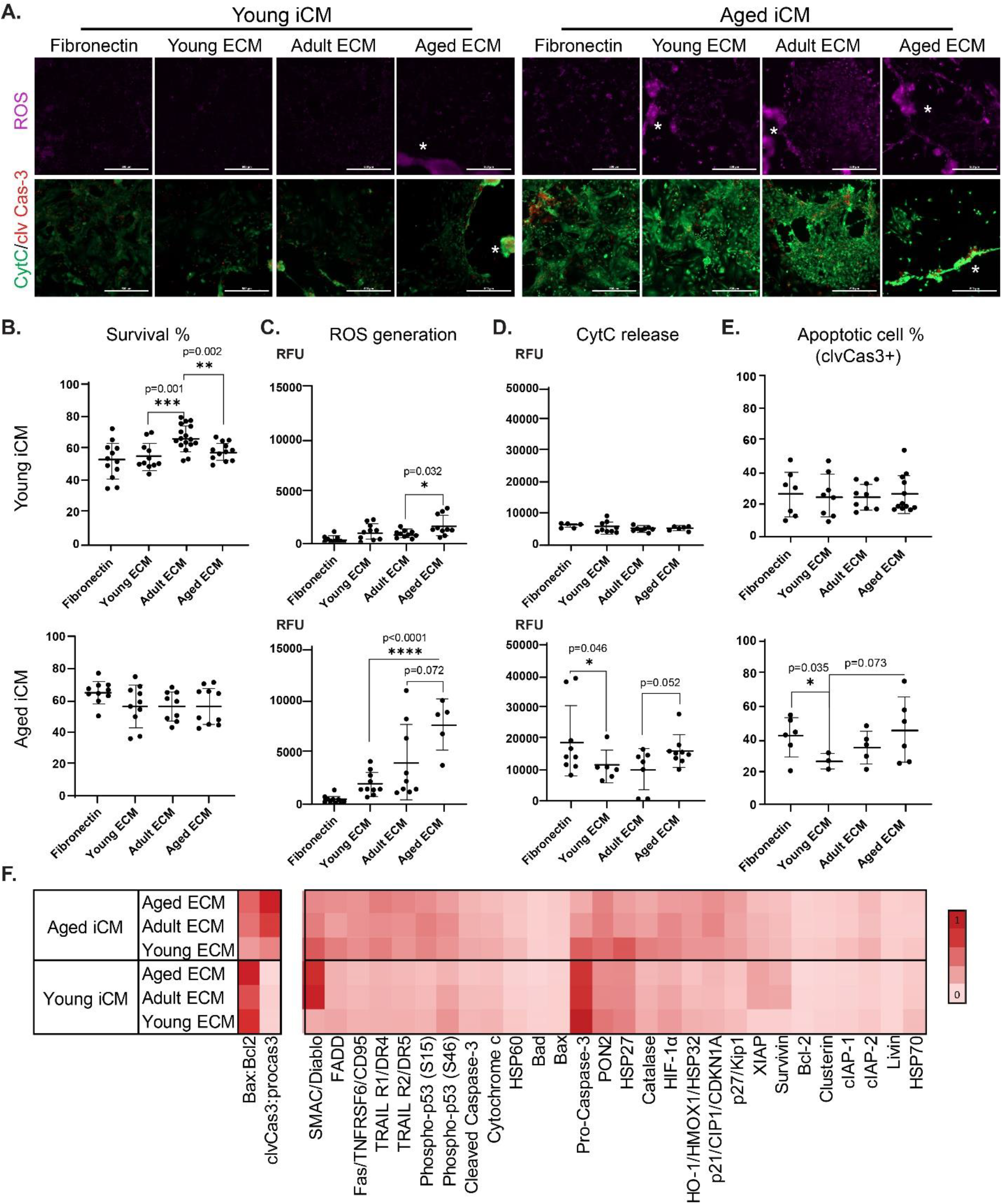
Cell survival and response assessment upon MI-mimicking stress conditions. (A) Apoptosis induction and execution related staining for ROS, Cytochrome-C and cleaved caspase-3 at 12 h-post RI (ROS-magenta, CytC-green, clvCas3-red). Scale bars represent 500μm. Detached cell clusters were shown with asterisk (white) on figures. Quantification of (B) cell survival rate calculated as live cell percentage, (C) ROS generation, (D) Cytochrome C release, and (E) apoptotic cell percentage calculated as clvCas3 expressing cell ratio. (F) Apoptosis-related protein expressions upon MI-mimicking stress conditions. Apoptotic rate at upstream and downstream of Cytochrome C release was calculated as Bax to Bcl-2 ratio and clvCas3 to proCas3 ratio, respectively. Statistical analysis was done using one-way ANOVA with post-hoc Tukey’s test. **** p<0.0001, ***p<0.001, **p<0.01, *p<0.05, n≥3. Data presented as mean ± standard deviation (SD).

We detected no ECM age-dependency in the young cell I/RI response. Aged cells, on the other hand, displayed an increasing CytC release and clvCas3 activity with increasing ECM age. For both young and aged cells, we noticed a large proportion of post-I/RI dead cells on the aged ECM **(Fig 5A)**.

Finally, cells were screened for apoptosis-related proteins, and the apoptotic rates at two stages were calculated: (i) the ratio of apoptosis regulators acting upstream of cytochrome C (Bax to Bcl-2), and (ii) the ratio of apoptosis executioner caspases (clvCas3 to pro-caspase 3). The upstream signals were abundantly expressed in the young cells, whereas the downstream signals were abundantly expressed in aged cells. Both apoptotic signals increased with increasing ECM age **(Fig. 5F)**. Proteins involved in the endogenous defense mechanism against oxidative stress, PON2, and HSP27 [72,73] were highly expressed in the young cells. Aged cells, on the other hand, expressed considerably more apoptosis-related proteins and their apoptotic rate followed an ECM age-dependent increase. Aged cells expressed high PON2, HSP27, and catalase on young ECM, which allowed them to cope with the elevated SMAC/Diablo. On aged ECM, aged cells expressed more death receptors TRAILR1 and TRAILR2[74] and less PON2, HSP27, HIF1a, and catalase **(Fig. 5F)**.

Overall, aged cells had impaired stress coping machinery and hence were more susceptible to MI-mimicking stress conditions. On young ECM, the endogenous defense mechanism was supported, and the severity of the stress-induced damage increased with increasing ECM age.

## 4. Discussion

We describe for the first time the individual and combined effects of cell and ECM aging on cardiomyocyte cell state, function, and response upon MI-mimicking stress conditions **(Fig. 6)**. We demonstrated that the young ECM held longevity, self-renewal, proliferation, and survival associated biochemical cues that promoted proliferation and drug responsiveness in young cells, additionally, induced cell cycle re-entry, and supported stress coping ability in the aged cells. We also showed that the adult ECM had biochemical cues supporting cardiac function that yielded faster calcium transient and mature ventricular type beating of the cells. Additionally, we found that the aged ECM presented aging cues that accelerated the aging phenotype and impaired cardiac function and stress defense machinery. Finally, we verified that the cardiac function declined with cellular aging and aged cells were more susceptible to stress conditions. Collectively, we provided a thorough analysis of cardiac aging to serve as a guide in the development of effective treatment strategies for CVDs with the inclusion of aged tissue microenvironment.

**Figure 6.**
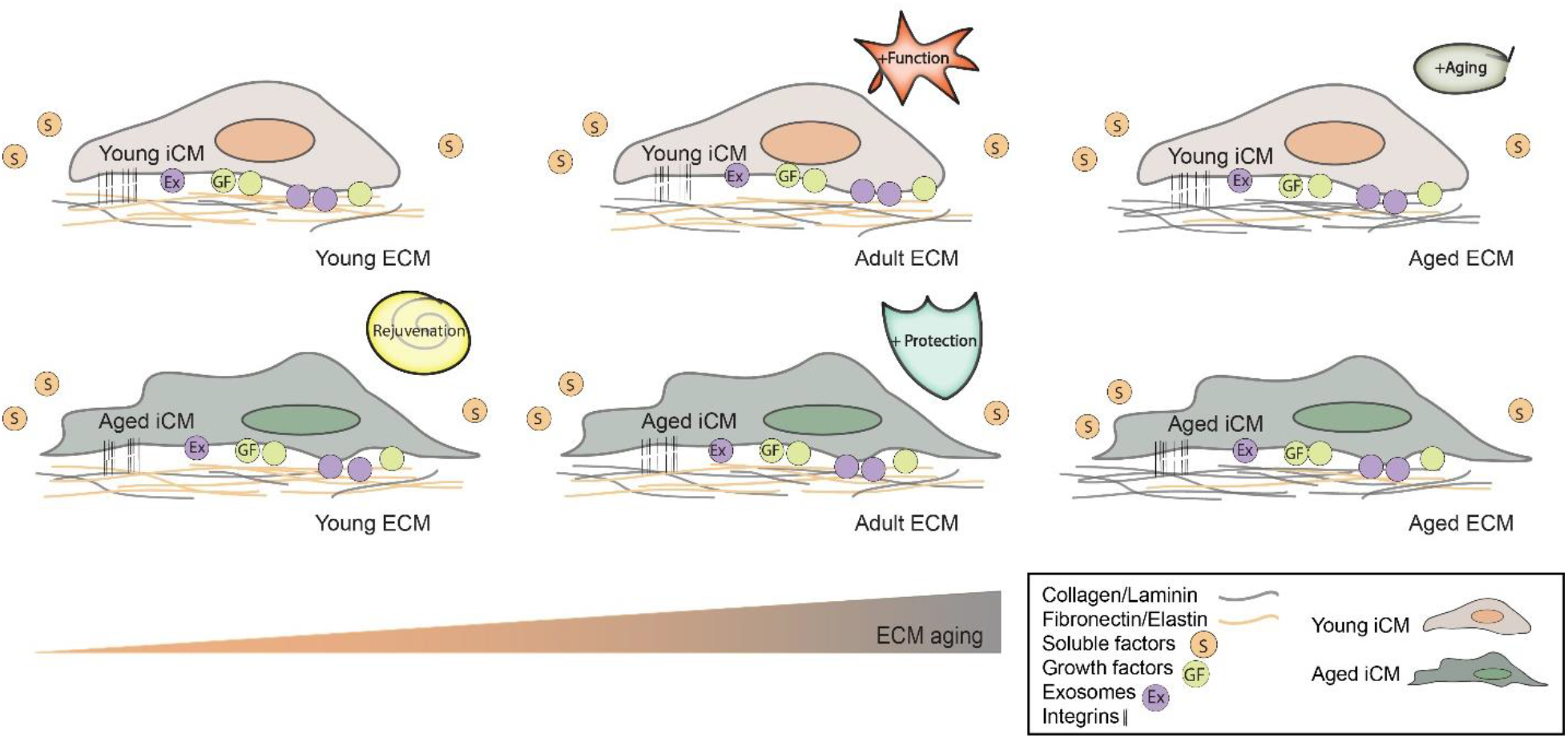
Study summary. Young iCMs (top) and aged iCMs (bottom) seeded on young, adult, and aged heart ECM. We detected compositional shifts with cardiac aging (showed with a color shift). ECM aging had different influence on the cardiac cell state, function, and response to MI-mimicking stress conditions. The major effects of the cardiac ECM age were i) adult ECM improved young cell cardiac function, ii) aged ECM fastened the aging process in the young cells,iii) young ECM rejuvenated aged cells, iv) adult ECM protected aged cells upon stress. Young cells on young ECM and aged cells on aged ECM served as controls.

Current regenerative therapies for heart diseases focus mainly on cellular aging and overlook the aged microenvironment surrounding the cells [75–81]. Although some groups have studied aging-associated biochemical changes in cardiac ECM [7,36], there is no information on the biochemical entities of advanced-age cardiac ECM and the interplay between different aged cells and environments reported to date. Here, we performed a thorough investigation of cardiac aging using human iPSC-derived cardiomyocytes and solubilized mouse heart ECMs recapitulating three age groups as young, adult, and aged. We evaluated key aspects of cell aging, cardiac maturity, and created MI-mimicking conditions to assess cell survival and response. This experimental design allowed us to dissect the effect of cellular and ECM aging and identify the biochemical changes that occur in the ECM, which will enhance the understanding of the basic biology of cardiac aging.

In our previous studies, we fully characterized iCMs differentiated from two separate human iPSC lines for their *in vitro* aging-related biomarkers as well as their maturity. We have demonstrated both cellular and functional deterioration and verified the loss of proliferative ability and poor calcium handling leading to impaired beating properties in cells cultured over 100 days^17^. In addition, in the current study, we discovered that some of the chronologically aged cells were not senescent but were residing in a quiescent state and re-entered the cell cycle in response to the extrinsic signals coming from the young ECM.

Here we observed that young cells were more responsive in terms of their ability to sense and respond to the environmental cues presented in the cardiac ECM. Conversely, aged cells exhibited a substantially decreased responsiveness to their environment. The desensitization of aged cells to changing ECM age can be explained by the impaired cellular activity and function, which is also seen in the aging human heart. The biochemical cues such as growth factors might not be reaching their ultimate targets in aged cells to alter the cell state or function [82,83]. Under MI-mimicking stress conditions, we observed cellular age-dependent increase of apoptosis rate. At the molecular level, unique expression patterns of apoptosis-related proteins reflected the influence of age on cell’s response to stress. Young cells had notably high stress coping elements such as protein chaperons and antioxidants **(Fig. 5F)**, that overruled the impact of the ECM age in stress response. Post-I/RI, we detected early apoptotic signals for young cells and late apoptotic signals for aged cells **(Fig. 5F)**. This was also verified with the endpoint results showing that aged cells were more inclined to activate stress associated pathways and apoptosis. These results suggested either a late onset of apoptosis or delayed progression of apoptosis in young cells. Overall, the observed differences between young and aged cells implied that prolonged culture is a simple yet useful way to obtain physiologically more relevant cell age groups. Our results showed that the ‘young cells’ could better mimic the young individuals whereas the ‘aged cells’ could better mimic the aged individuals; and this information is valuable considering most of the studies have been conducted using the young cells and the main risk group for the heart diseases is the older individuals.

Additionally, here we recorded distinct responses and activated pathways between young and aged cells both in health and disease conditions. Cellular state **(Fig. 3)** and cardiac function **(Fig. 4)** were considerably altered by the ECM age, emphasizing the importance of age-associated biochemical changes occur in ECM and the potential use of ECM as a therapeutic tool.

In order to study the effect of the ECM age on cardiac function and stress response, we used young, adult, and aged mouse cardiac ECM. We verified the preservation of the main ECM biochemical entities in the decellularized cardiac tissues. Similar to other studies[7,78], we detected compositional shifts in decellularized ECMs with aging **(Fig. 2)** and extended the literature knowledge by providing information on the advanced age (22-24 month-old) mouse cardiac ECM which has not been reported before. Interestingly, the influence of the solubilized cardiac ECM as a coating material sustained up to 3 weeks in culture, although the seeded iCMs on these coated surfaces also deposited a considerable amount of new ECM within 3 days of culture, yet to different degrees depending on the age state of the cell and the ECM **(Supplementary Fig. 4)**. This suggested the dynamic interplay between the cells and the ECM: (i) the ECM coating influenced *de novo* ECM secretion by the cells and (ii) cells respond to that resulting deposited-coated ECM mixture.

Mimicking the early developmental stages of the heart, young ECM had notably high elastic matrix and proliferation-associated cues **(Fig. 2C)**. As previously reported in fetal hearts[84], young ECM promoted young cell proliferation **(Figs. 3B and 2C)**. Surprisingly, young ECM also induced cell cycle reentry of aged cells that we have considered senescent due to increased senescence markers and lost proliferative abilities **(Fig. 3C)**. Recent studies reported that only a small number of cardiomyocytes can complete mitosis at an adult stage, and there are many conflicting results on the evaluation of karyokinesis and cytokinesis of these cells[76,85–88]. Here, we detected some aged iCMs undergoing mitosis after a 3-week incubation on the young ECM **(Fig. 3B and Supplementary Fig. 2)**. This suggested the biochemical composition of the cardiac ECM alone is a potent agent to alter the cell state in such a short treatment period. Besides regeneration, our results suggested the pro-survival cues in young ECM might be the driving force behind the increased production of protein chaperons and antioxidants in aged cells **(Figs. 1C, 3F, and 5F)**. Hence, we concluded that young ECM supported aged cell defense machinery and resulted in lower ROS generation and apoptotic activation **(Fig. 5)**.

Although previous studies offered the use of ECM or ECM-based materials for cardiac regeneration and repair [28,89–91], our results highlight the importance of the ‘age’. *In situ* delivery of young cardiac ECM holds the potential to stimulate the self-renewal abilities of the patient’s resident cells in the heart and aid cells in coping with oxidative stress conditions. The beating frequencies of iCMs have been reported below the human adult cardiomyocyte beating rate (∼1 Hz [92]) as 0.3-0.6 Hz[14,93,94]. Here, only after a 3-week-of-culture on adult ECM, young iCM displayed robust tissue-like beating at a rate of 0.80±0.15 Hz **(Figs. 4A-B)**. Our results demonstrated that the adult ECM holds the potential to improve the cardiac function in both healthy and post-MI heart. The cardiac homeostasis and function related cues detected in the adult ECM might explain the faster calcium transient and facilitated beating observed **(Fig. 4)**. In accordance with our results, structural and functional protection properties of the adult ECM have also been reported in brain [95]. Currently, cell therapies often result in minimal functional improvement in the post-MI heart [14–17]. This is mostly because *in vitro* cultures lack the ability to fully mimic *in vivo* cardiac maturation [18] and even after biochemical and electromechanical stimulations iCMs display structural and functional immaturity. The adult ECM supplement in cell therapy holds the potential to improve the electromechanical coupling of the delivered cells to the host cardiac cells and prevent arrhythmias. Alternatively, adult ECM delivery to the patient hearts with minor functional defects might rewire the calcium handling and affect beating kinetics to match the healthy cardiac cells in a post-MI heart.

Finally, our results showed that aged ECM enhanced cardiac structural maturity **(Figs. 3B and 3E)**. However, it adversely affected the cardiac function **(Fig. 4)** by causing longer than functionally acceptable sarcomeres and potential hypertrophy. We have detected high amounts of profibrotic and hypertrophy-associated cues in the aged ECM that might also promote myofibrillar remodeling[96]. Therefore, due to its adverse effects, the use of aged ECM in the clinic should not be considered. The alternative use of the aged ECM could be to recapitulate an aging heart environment in *in vitro* disease modeling studies to obtain physiologically more relevant responses.

## Conclusion

In conclusion, we show that the cell’s response to the ECM age depends highly on the cell age, and aging dramatically reduces cell responsiveness. We also show that ECM age influences the proliferative ability, maturity, and stress response of both young and aged cells. These observations highlight the importance of cellular and environmental age in a patient’s response to treatment. Delivering (i) young ECM to aged heart holds the potential to induce regeneration, and (ii) a mixture of young iCMs and adult ECM to aged heart holds potential to improve cardiac function. Since the main risk group is older patients, better understanding the individual and combinatorial effects of different age components is highly useful for the patient and complication-specific treatments.

## Acknowledgements

This work was funded by the National Science Foundation CBET grant number [1651385] and [NSF-1805157]. We would also like to thank Dr. Sharon Stack and her lab for providing mice tissues.

## Data availability statement

The raw/processed data required to reproduce these findings can be shared upon request.

## Author Contribution

**S. Gulberk Ozcebe:** Conceptualization, Methodology, Investigation, Data curation, Visualization, Formal analysis, Writing-Original draft preparation, Writing - Review & Editing **Gokhan Bahcecioglu:** Formal analysis, Writing-Reviewing and Editing **Xiaoshan S. Yue:** Data curation **Pinar Zorlutuna:** Supervision, Conceptualization, Methodology, Formal analysis, Funding acquisition, Resources

---

**Supplementary Figure 1.**
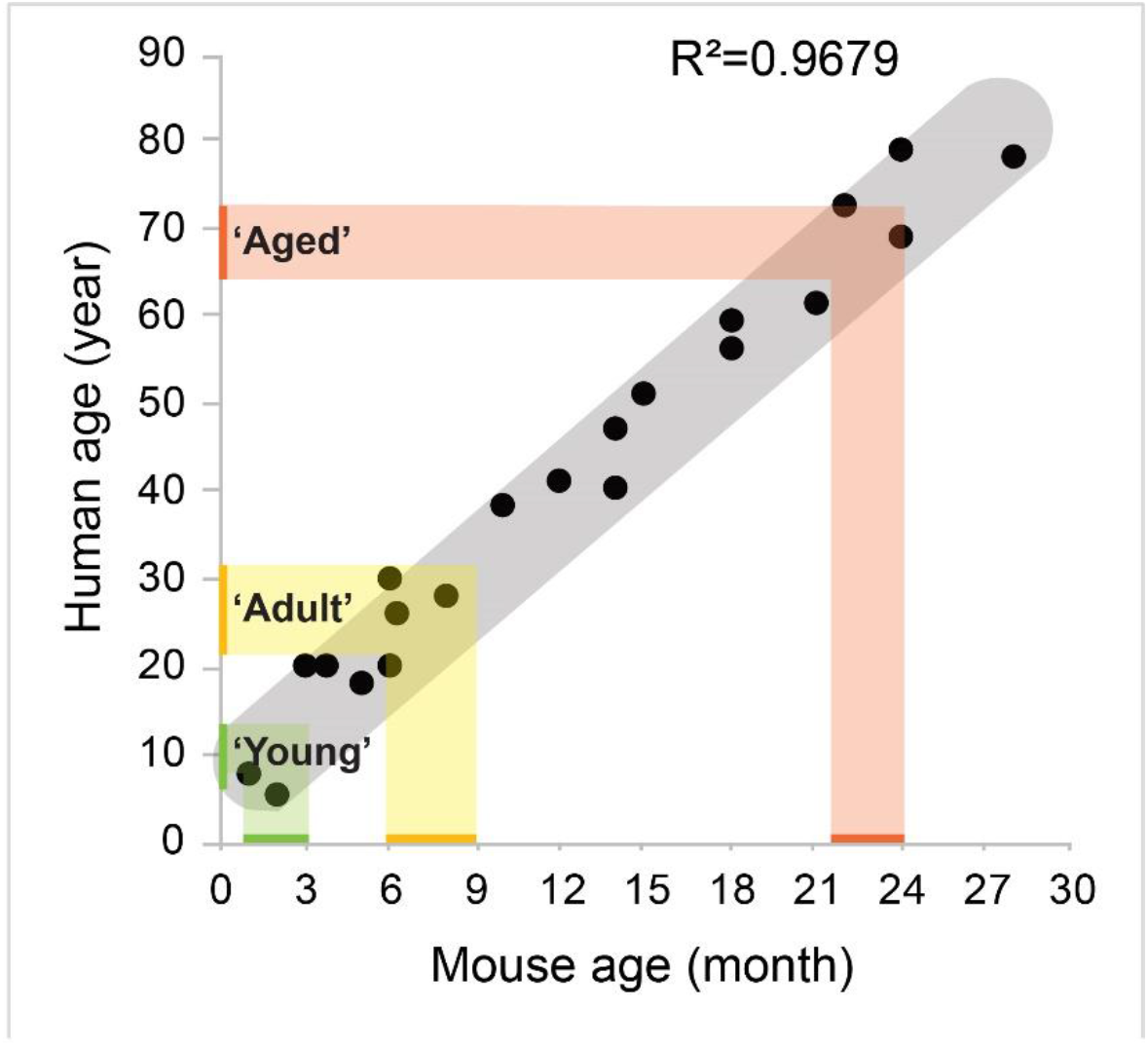
Correspondence between human and C57BL/6J mice lifespans. Data collected from previous studies [38], [40], [42] to compare mouse age to human age. Between 3-27-months, the mouse and human age correlation is almost linear (R=0.97). The age range of the young (1-3-months, green), adult (6-9-months, yellow) and aged (22-24-months, orange) mice used in our study is indicated. The linear fit estimates the human age corresponding to the indicated mouse ages as 10-16 years, 24-32 years, and 67-73 years.

**Supplementary Figure 2.**
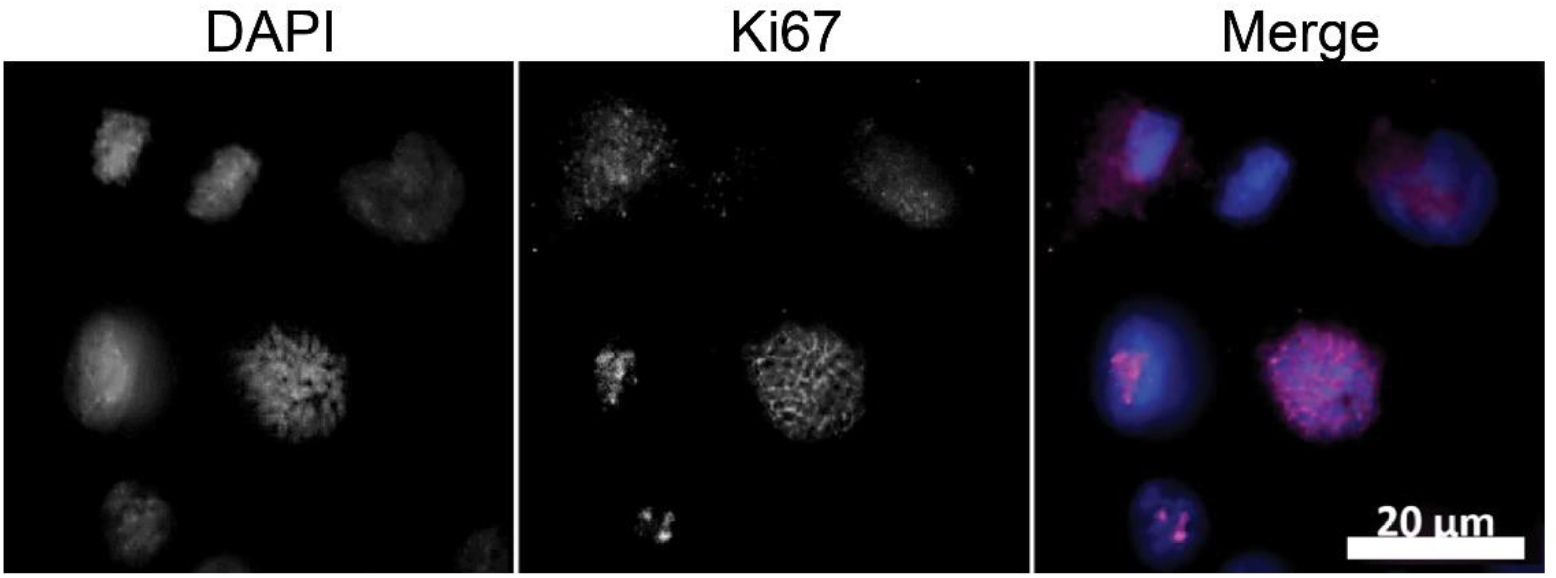
Expression of Ki67 in aged cells cultured on young ECM. Aged cells undergoing mitosis and karyokinesis captured. (DAPI-blue, Ki67-magenta)

**Supplementary Figure 3.**
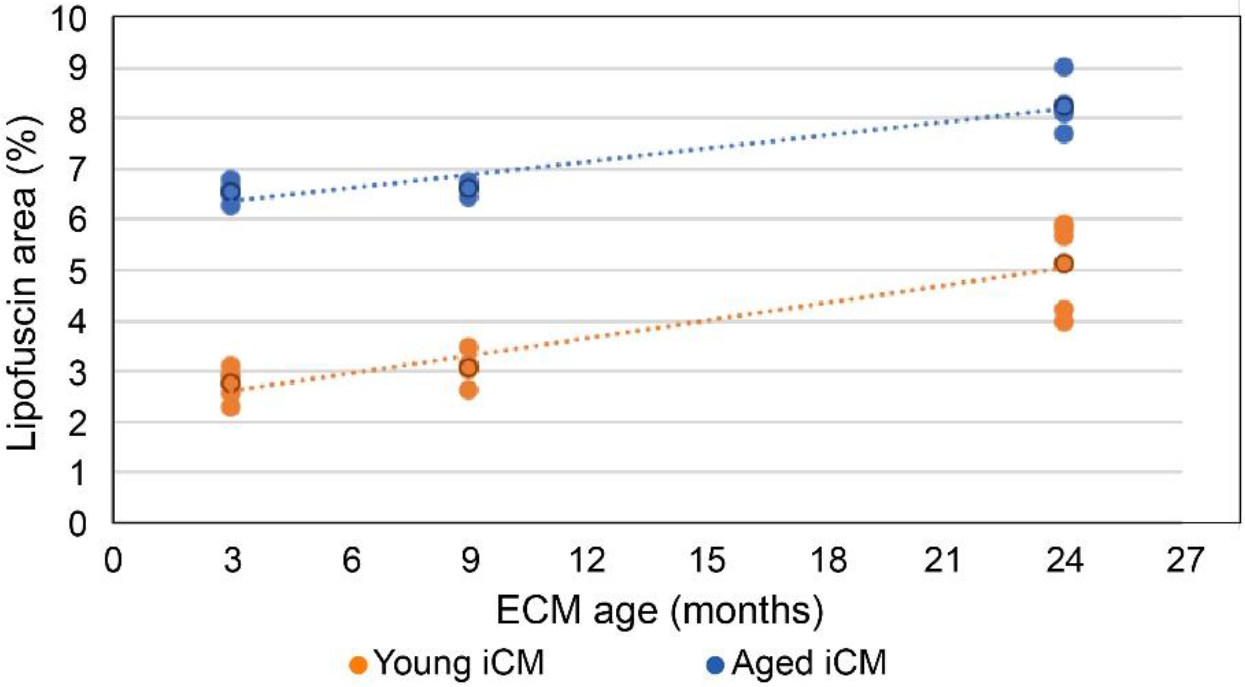
Linear correlation between ECM age and the lipofuscin accumulation. Lipofuscin accumulation (%) of the young iCM (orange) and aged iCM (blue) show linear increase with ECM aging.

**Supplementary Figure 4.**
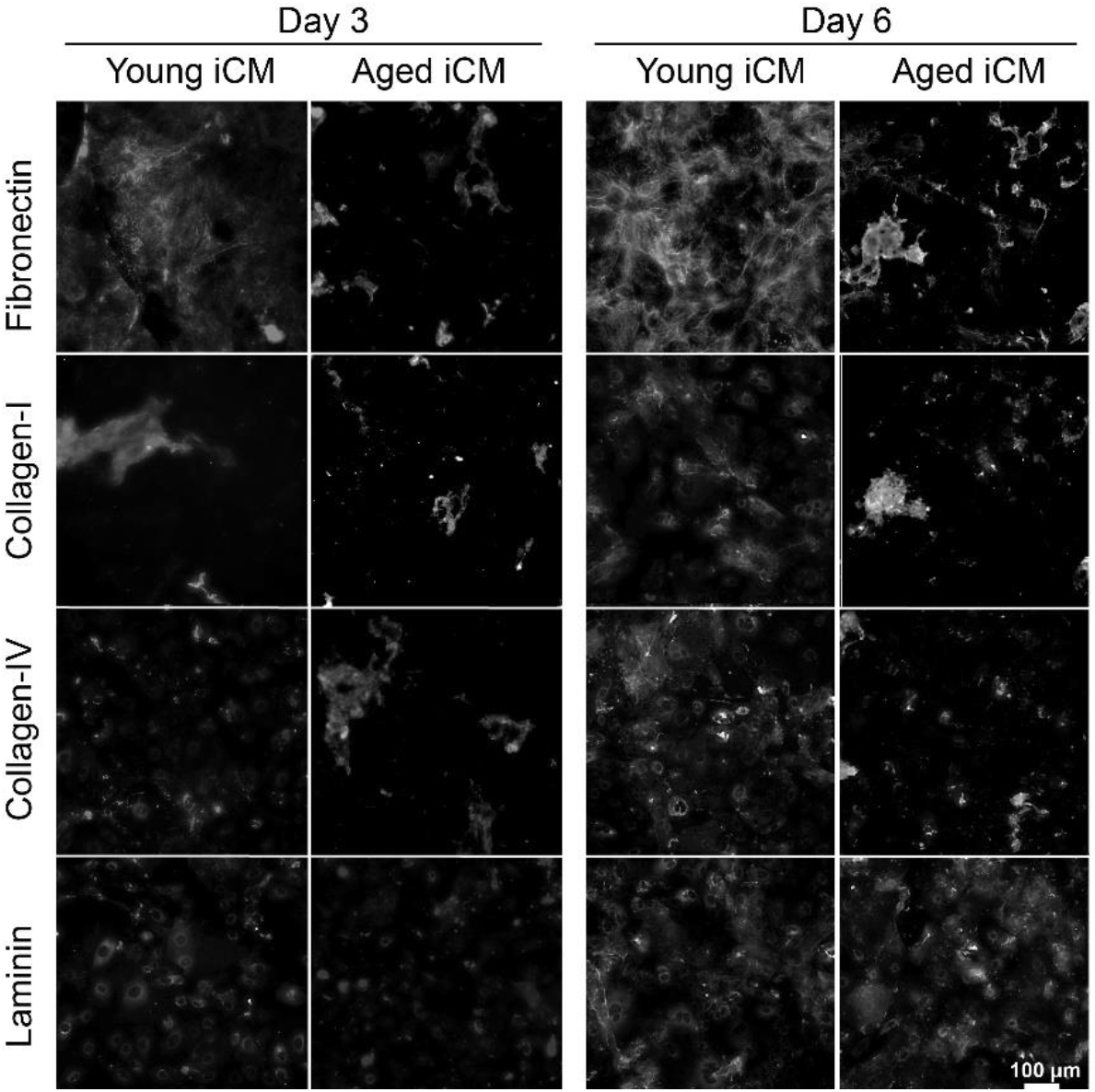
ECM deposition by the iCMs. Immunostaining of the common ECM proteins, Fibronectin, Collagen-I, Collagen-IV and Laminin deposited by the young and aged iCMs on day 3 (left panel) and day 6 (right panel).

